# Enzymatic Specificity of Conserved Rho GTPase Deamidases Promotes Invasion of *Vibrio parahaemolyticus* at the Expense of Infection

**DOI:** 10.1101/2022.06.13.496033

**Authors:** Alexander E. Lafrance, Suneeta Chimalapati, Nalleli Garcia Rodriguez, Lisa N. Kinch, Karan Gautam Kaval, Kim Orth

## Abstract

*Vibrio parahaemolyticus* is among the leading causes of bacterial seafood-borne acute gastroenteritis. Like many intracellular pathogens, *V. parahaemolyticus* invades host cells during infection by deamidating host small Rho GTPases. The Rho GTPase deamidating activity of VopC, a type three secretion system (T3SS) translocated effector, drives *V. parahaemolyticus* invasion. The intracellular pathogen uropathogenic *Escherichia coli* (UPEC) invades host cells by secreting a VopC homolog, the secreted toxin cytotoxic necrotizing factor one (CNF1). Because of the homology between VopC and CNF1, we hypothesized topical application of CNF1 during *V. parahaemolyticus* infection could supplement VopC activity. Here, we demonstrate that CNF1 improves the efficiency of *V. parahaemolyticus* invasion, a bottleneck in *V. parahaemolyticus* infection, across a range of doses. CNF1 increases *V. parahaemolyticus* invasion independent of both VopC and the T3SS altogether, but leaves a disproportionate fraction of intracellular bacteria unable to escape the endosome and complete their infection cycle. This phenomenon holds true in the presence or absence of VopC, but is particularly pronounced in the absence of a T3SS. The native VopC, by contrast, promotes a far less efficient invasion, but permits the majority of internalized bacteria to escape the endosome and complete their infection cycle. These studies highlight the significance of enzymatic specificity during infection, as virulence factors (VopC and CNF1 in this instance) with similarities in function (bacterial uptake), catalytic activity (deamidation), and substrates (Rho GTPases) are not sufficiently interchangeable for mediating a successful invasion for neighboring bacterial pathogens.

**IMPORTANCE:** Many species of intracellular bacterial pathogens target host small Rho-GTPases to initiate invasion, including the human pathogens *Vibrio parahaemolyticus* and uropathogenic *Eschericia coli* (UPEC). The type three secretion system (T3SS) effector VopC of *V. parahaemolyticus* promotes invasion through the deamidation of Rac1 and CDC42 in the host, whereas the secreted toxin cytotoxic necrotizing factor one (CNF1) drives UPEC’s internalization through the deamidation of Rac1, CDC42, and RhoA. Despite these similarities in the catalytic activity of CNF1 and VopC, we observed the two enzymes were not interchangeable. Although CNF1 increased *V. parahaemolyticus* endosomal invasion, most intracellular *V. parahaemolyticus* aborted their infection cycle and remained trapped in endosomes. Our findings illuminate how the precise biochemical fine-tuning of T3SS effectors is essential for efficacious pathogenesis. They moreover pave the way for future investigations into the biochemical mechanisms underpinning *V. parahaemolyticus* endosomal escape, and more broadly, the regulation of successful pathogenesis.

## Introduction

Bacterial pathogens are masters of biochemistry, utilizing a diverse array of virulence factors to acquire nutrients from the host, avoid the host immune response, and modulate host signaling during infection (1). Many species of Gram-negative bacteria couple these strategies with invasion of host cells to establish an intracellular replicative niche. Factors that mediate bacterial invasion of the host may take many forms, but their mechanisms of action commonly converge on Rho GTPase signaling within host cells (2). Rho GTPases act as molecular switches that are active when bound to their GTP substrate, enabling them to interact with downstream signaling proteins. Once a Rho GTPase hydrolyzes GTP to GDP, however, the enzyme undergoes a conformational change and can no longer interact with its downstream targets (3). Three of the best-characterized Rho GTPases in eukaryotic cells are RhoA, Rac1, and CDC42, which contribute to the formation of stress fibers, filipodia, and lamellipodia, respectively, through the regulation of actin polymerization (3).

As regulators of myriad downstream processes in the cell and of one another, Rac1, RhoA, and CDC42 represent prime targets for manipulation by intracellular pathogens during the host invasion process. *Vibrio parahaemolyticus*, a Gram-negative, halophilic bacterium endemic to estuarine and marine environments, is one such organism that avails itself of these targets. *V. parahaemolyticus* possesses a wide variety of virulence factors, including two type three secretion systems (T3SS) and secreted hemolysins. Most relevant to this study is the second T3SS, T3SS2, that mediates a Rac1- and CDC42-dependent invasion of the host (4–6). The effector CNF1-family deamidase VopC constitutively activates Rac1 and CDC42 through the deamidation of the conserved glutamine 61 residue on both proteins, inducing actin polymerization at the plasma membrane, membrane ruffling, and finally, uptake of the bacteria into the host (Fig. 1A) (4, 7). VopC translocation, and thus its deamidase activity, is contingent on the presence of an N-terminal T3SS secretion signal and chaperone binding domain (Fig. S1A) (8). Once inside, *V. parahaemolyticus* inhibits MAPK signaling in the host, stimulates stress fiber formation and bundling, cripples the host reactive oxygen species response, and induces cytotoxicity through multiple mechanisms in addition to enterotoxicity (9–14).

**Fig 1.**
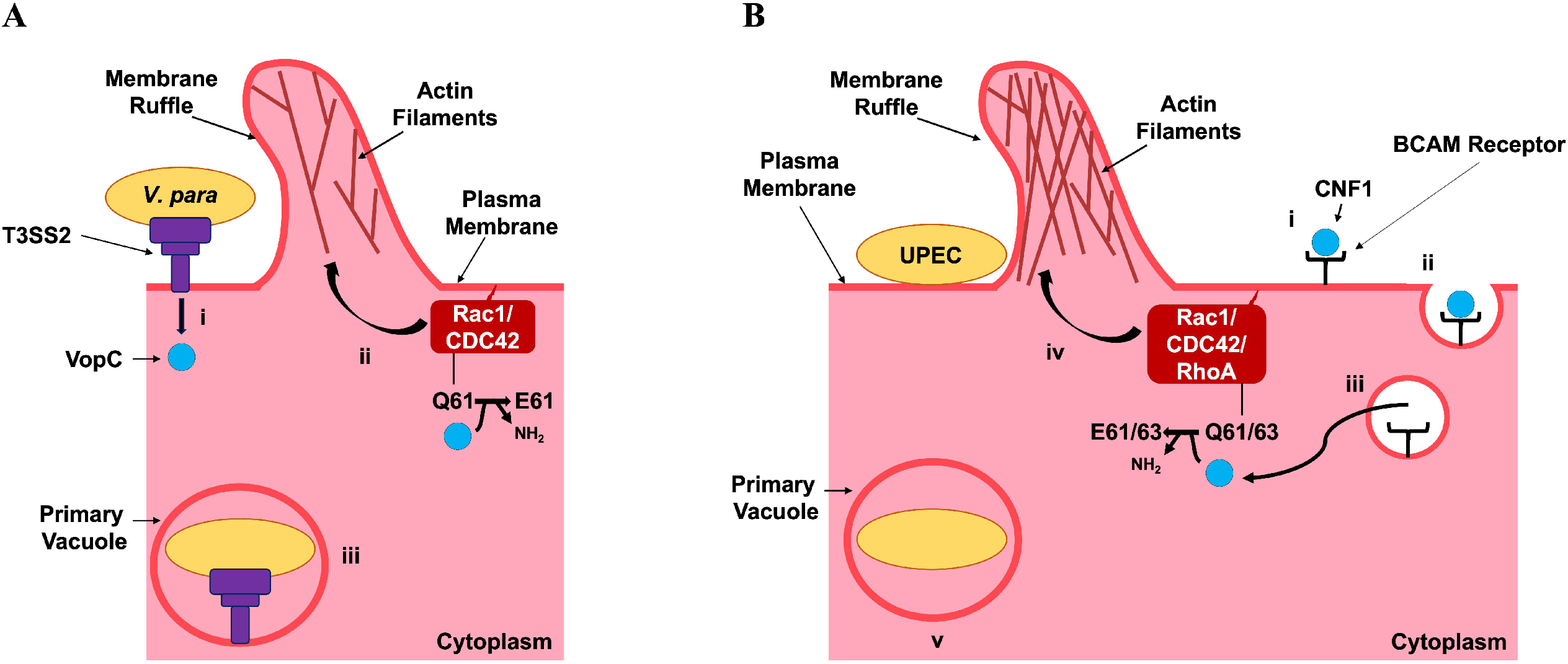
Comparison of the invasion mechanisms of *V. parahaemolyticus* and UPEC. **(A)** *V. parahaemolyticus* (i) adheres to the host cell surface, where it translocates effectors across the host cell membrane into the cytoplasm through the T3SS2, including VopC. (ii) VopC deamidates glutamine 61 of both Rac1 and CDC42 at the plasma membrane, constitutively activating the Rho GTPases to promote the polymerization of actin branches and ruffling of the plasma membrane. (iii) Membrane ruffles engulf *V. parahaemolyticus*, culminating in its internalization in a primary vacuole. **(B)** UPEC (i) adheres to the host cell surface and secretes the toxin CNF1 into the extracellular space through an unknown mechanism. CNF1 then binds Lutheran basal cell adhesion molecule (BCAM) receptors. (ii) CNF1 is endocytosed. (iii) acidification triggers CNF1 cleavage and export through an unknown mechanism into the host cytoplasm. (iv) CNF1 deamidates glutamine 61 of Rac1 and CDC42, and glutamine 63 on RhoA, constitutively activating all three Rho GTPases to promote actin filament polymerization and ruffling of the plasma membrane. (v) Membrane ruffles engulf UPEC, culminating in its internalization in a primary vacuole.

*V. parahaemolyticus* is not the only bacteria to utilize complex secretion systems to target Rho GTPases during infection. For example, the T3SS effector Pnf of insect pathogen *Photorhabdus asymbiotica* both deamidates and transglutaminates RhoA and Rac1 to facilitate invasion (15). The enteric pathogen *Salmonella enterica* also translocates multiple effectors into the host through a T3SS, including SopE, SopE2, and SopB, to stimulate invasion through CDC42 and Rac1 activation. However, these effectors function as exchange factors (GEFs) that activate CDC42 and Rac1 by charging them with a steady supply of GTP substrate, rather than constitutively activating the Rho GTPases through deamidation or another modification (16). Interestingly, *Burkholderia cenocepacia*, an opportunistic pathogen commonly associated with lung infections in human cystic fibrosis patients, delivers TecA, its own deamidase of RhoA, Rac1, and CDC42, into the host cytoplasm by way of a toxin-loaded “harpoon” called a type six secretion system (T6SS) (17). Unlike VopC and Pnf, however, the deamidation of Rho GTPase residues targeted by TecA result in their deactivation, not the constitutive activation (17).

Like *V. parahaemolyticus*, uropathogenic *Escherica coli* (UPEC) also mediates an intracellular infection by deamidating host Rho GTPases with the toxin cytotoxic necrotizing factor one (CNF1). Unlike *V. parahaemolyticus*’s VopC, however, CNF1 targets RhoA, in addition to CDC42 and Rac1, and is secreted into the extracellular space rather than translocated through a T3SS (18). CNF1 contains five primary domains (Fig. S1A). The first three domains (D1-3) cumulatively responsible for interacting with the host p37LR laminin receptor precursor protein and translocation into the host (19, 20). The latter two domains of CNF1 (D4-5) respectively encode an ADP-ribosyltransferase-like domain of unknown function, DUF4765, and a deamidase domain fused to a Lutheran (Lu) adhesion glycoprotein/ basal cell adhesion molecule (BCAM) binding motif (19–21). The exact manner by which CNF1 is secreted remains unknown, as it does not appear to contain a cleavable secretion signal; however, interaction with both p37LR and Lu/BCAM receptors appear to be necessary to trigger toxin endocytosis after secretion (7, 19–22). Within the vacuole CNF1 is activated by pH-dependent cleavage, and subsequently enters the host cytoplasm through an unidentified mechanism, where it deamidates Rac1, RhoA, and CDC42 at the plasma membrane to induce membrane ruffling and promote bacterial internalization (Fig. 1B) (21, 23).

Many different bacteria have adopted invasion strategies combining the Rho GTPase deamidation invasion mechanism of the CNF superfamily with a secreted cytotoxin delivery mechanism (24, 25). For example, an isolate of calf and piglet pathogenic *E. coli* was found to contain a cytotoxin deemed CNF2 with 90% homology to CNF1, and *Yersinia pseudotuberculosis* secretes the cytotoxin CNFy, which bears a 65.1% homology to CNF1 (24, 26). Multiple *Bordetella* species also carry the DNT (dermonectrotizing toxin), which has been shown to activate RhoA, Rac1, and CDC42 by transglutaminating the same residues deamidated by CNF1 and its aforementioned homologues; however, DNT does not appear to be secreted, or even to leave the bacterial cytoplasm during infection (24, 27).

Despite the considerable homology between these conserved Rho GTPase deamidases, differences in domain organization and sequences across these enzymes contribute to substantial diversity among them. Evolutionary pressure modulates the catalytic activity, efficiency, and temporal regulation of otherwise similar effectors to operate efficiently in concert with biochemical arsenals unique to each bacterial species (28). Such differences between homologs are thus highly illustrative of the differences between life cycles and infection mechanisms of bacterial pathogens. Moreover, although individual variations between these homologs may prove challenging to identify, aggregate differences between homologs such as those within the CNF deamidase superfamily can be identified by testing the cross-compatibility of conserved virulence factors. As members of the CNF1 deamidase superfamily, the catalytic domains of VopC and CNF1 share only a 24% amino acid sequence identity, but contain conserved catalytic cysteine and histidine residues (Fig S1) (4). We consequently hypothesized, based on the enzymes’ homology, that CNF1 could complement VopC in mediating bacterial invasion of the host (Fig. S1) (5, 24, 29).

To test our hypothesis, we began with the *V. parahaemolyticus* CAB2 strain, derived from the clinical isolate RIMD2210633, in which the hemolysin-encoding genes and the transcriptional regulator of the first T3SS have been deleted (Table 1) (4). With this strain, we were able to assess the importance of the T3SS2 and its effectors during *in vitro* infection assays without interference by other virulence systems. To determine whether CNF1 could promote invasion of *V. parahaemolyticus*, purified CNF1 was added to HeLa cells during infection with CAB2. We observed that CNF1 correlated with increased invasion of CAB2 across a range of doses, and that CNF1 treatment did not adversely affect HeLa cells. We also found that treatment of HeLa cells with a catalytically inactive mutant of CNF1 during infection with CAB2 did not significantly impact bacterial invasion. Next, we assessed the effect of CNF1 treatment on invasion-deficient mutants CAB2Δ*vopC*, which does not express the T3SS2 effector VopC necessary for invasion, and CAB4, which cannot express hemolysins, the T3SS1, or the T3SS2 (4). Neither strain was able to invade host cells after treatment with the catalytic dead mutant of CNF1; however, treatment with wild type CNF1 promoted the invasion of both CAB2Δ*vopC* and CAB4. CNF1 treatment, but not with the catalytic dead mutant of CNF1, increased *V. parahaemolyticus* invasion across all three strains. Although, few intracellular CAB2 and CAB2Δ*vopC* completed their infection cycle after internalization and the infections of nearly all intracellular CAB4 (Table1) were arrested under the same conditions. A closer examination of CNF1-treated infections revealed that aborted infections were marked by the confinement of intracellular bacterial to the endosome, while the minority of intracellular *V. parahaemolyticus* which did escape into the cytosol appeared capable of proliferating, and ultimately egressing from the host as normal. Ultimately, we observed the similarities between CNF1 and VopC were sufficient for CNF1 to drive *V. parahaemolyticus* invasion independent of VopC, but were insufficient for *V. parahaemolyticus* to establish a productive, intracellular replicative niche. These findings highlight the precision in biochemical regulation and signaling for bacterial pathogens and emphasize more broadly the importance of enzymatic specificity and delivery in mediating the tightly-orchestrated processes of bacterial pathogenesis.

**Table 1.** Summary of *V. parahaemolyticus* strains utilized in this study.

| Strain | Genotype | Description | Source |
| --- | --- | --- | --- |
| CAB2 | *Δtdh, Δtrh, ΔexsA* | Deleted TDH & TRH hemolysins, deleted T3SS1 transcriptional regulator | (60) |
| CAB2-GFP | *Δtdh, Δtrh, ΔexsA* +pGFP | Deleted TDH & TRH hemolysins, deleted T3SS1 transcriptional regulator, transformed with pGFP | (31) |
| CAB2ΔvopC | *Δtdh, Δtrh, ΔexsA, ΔvopC* | Deleted TDH & TRH hemolysins, deleted T3SS1 transcriptional regulator, deleted T3SS2 effector VopC | (4) |
| CAB2ΔvopC-GFP | *Δtdh, Δtrh, ΔexsA, ΔvopC*, +pGFP | Deleted TDH & TRH hemolysins, deleted T3SS1 transcriptional regulator, deleted T3SS2 effector VopC, transformed with pGFP | Unpublished (Marcela de Souza Santos) |
| CAB4 | *Δtdh, Δtrh, ΔexsA, ΔvtrA* | Deleted TDH & TRH hemolysins, deleted T3SS1 & T3SS2 transcriptional regulators | (60) |
| CAB4-GFP | *Δtdh, Δtrh, ΔexsA, ΔvtrA*, +pGFP | Deleted TDH & TRH hemolysins, deleted T3SS1 & T3SS2 transcriptional regulators, transformed with pGFP | Unpublished (Marcela de Souza Santos) |

## Results

### Purified CNF1 improves the invasion efficiency of CAB2

To determine the capacity of CNF1 to promote bacterial invasion in HeLa cells, we first expressed and purified an N-terminally 6xHis-tagged CNF1 using standard biochemical techniques (Fig. S2) (30). We subsequently performed a gentamicin protection assay, in which CAB2, that had been induced with bile acids to express the T3SS2, was allotted 2 hours to invade HeLa cells in the presence of 0, 0.5, 2.5, or 10 µg/mL CNF1 before the application of gentamicin, which killed all extracellular bacteria and spared the intracellular bacteria for quantification (31). Consistent with previous research, addition of CNF1 (up to 10 µg/mL) did not adversely affect the health of HeLa cells after the 2 hour treatment (32). The surviving intracellular bacteria were quantified at 1 hour post-gentamicin treatment (PGT), when the bacteria had invaded and begun replicating within the endosome (31). CNF1 at all three tested concentrations correlated with a significant increase in intracellular bacteria at 1 hour PGT, as evidenced by the elevated CFU counts relative to the untreated control. (Fig. 2A). These quantitative findings were corroborated qualitatively by confocal micrographs of HeLa cells infected with bile acid -induced, GFP-expressing CAB2 (CAB2-GFP) taken at 1 hour PGT. Noticeably, the number of cells invaded was higher as more GFP positive bacteria were observed inside a larger number of HeLa cells after 0.5, 2.5, and 10 µg/mL CNF1 treatment than in the untreated control (Fig 2B) (4). While no significant difference in bacterial invasion was observed between the different concentrations of CNF1, 2.5µg/mL CNF1 yielded the greatest amount of invasion on average, and therefore, was used for all future infection experiments (Fig. 2A).

**Fig 2.**
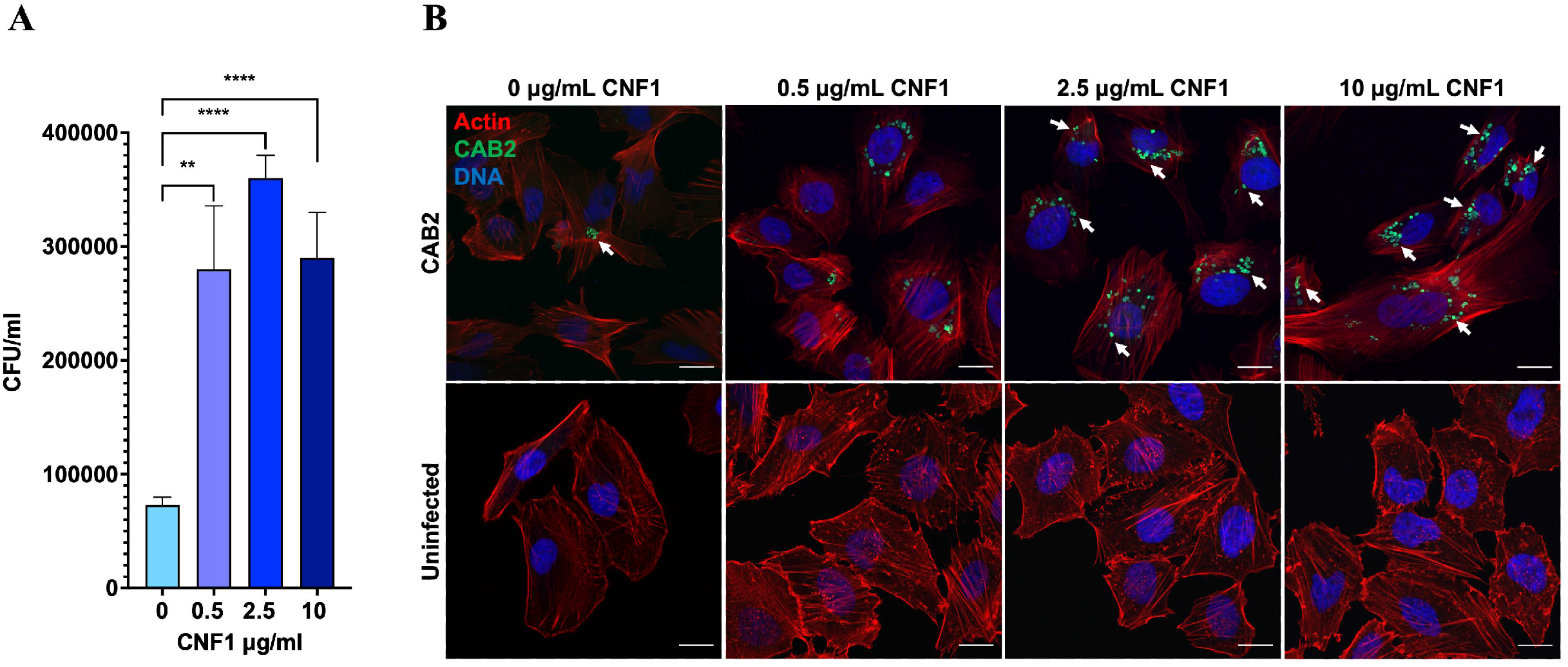
Supplementation of infection medium with CNF1 promotes invasion of CAB2 *V. parahaemolyticus*. **(A)** Gentamicin protection assay with CAB2 at an MOI = 10 demonstrates infection media supplemented with 0.5μg/mL, 2.5μg/mL, and 10μg/mL CNF1 2 hours prior to gentamicin application, and concurrent with the beginning of infection. Error bars represent standard deviation of three technical replicates. Statistical significance measured using one-way ANOVA and multiple comparisons test (***, P < 0.0005; ****, P < 0.00005). The gentamicin protection assay quantification shown here is a representative figure of three biological replicates. **(B)** Representative confocal micrograph of HeLa cells infected for 2 hours with CAB2-GFP at an MOI = 10 and treated with CNF1at a concentration of 0.5μg/mL, 2.5μg/mL, and 10μg/mL CNF1 for 2 hours (top). Uninfected control HeLa cell treated with CNF1 for 2 hours (bottom). All HeLa cells stained for F-actin using rhodamine-phalloidin (red) and Hoechst DNA stain (blue). White arrows indicate internalized CAB2-GFP. Scale bar, 20μm.

### CNF1 catalytic activity drives CAB2 invasion, culminating in aborted infections

Having established both an ideal working concentration of CNF1 and a correlation between CNF1 and increased invasion of HeLa cells by CAB2, we sought to test if the catalytic activity of CNF1 drove increased CAB2 invasion and to assess the impact of CNF1-mediated invasion on later stages of infection. We generated a catalytic dead CNF1 mutant as a negative control using PCR mutagenesis to replace the catalytic cysteine 866 residue with an alanine (CNF1 C866A). We then expressed and purified CNF1 C866A using the same standard biochemical techniques employed for the purification of wild type CNF1 (Fig. S2D) (33, 34).

We repeated the gentamicin protection assay and confocal microscopy analysis as described above but extended the gentamicin treatment time to 7 hours and collected intracellular CFU counts and confocal images at 1, 4, and 7 hours PGT. These time points were intended to capture CAB2’s behavior at the three major stages of infection of the canonical CAB2 infection cycle: 1) confinement to the endosome 2) active cytoplasmic replication and 3) egress from the host cell (31). Consistent with our earlier findings, CAB2 invasion was substantially increased in the presence of 2.5µg/mL CNF1 at 1 hour PGT based on CFU counts (Fig. 2A, 3A). Furthermore, treating HeLa cells with 2.5µg/mL CNF1 C866A did not increase invasion relative to the untreated control (Fig. 2A, 3A).

**Fig 3.**
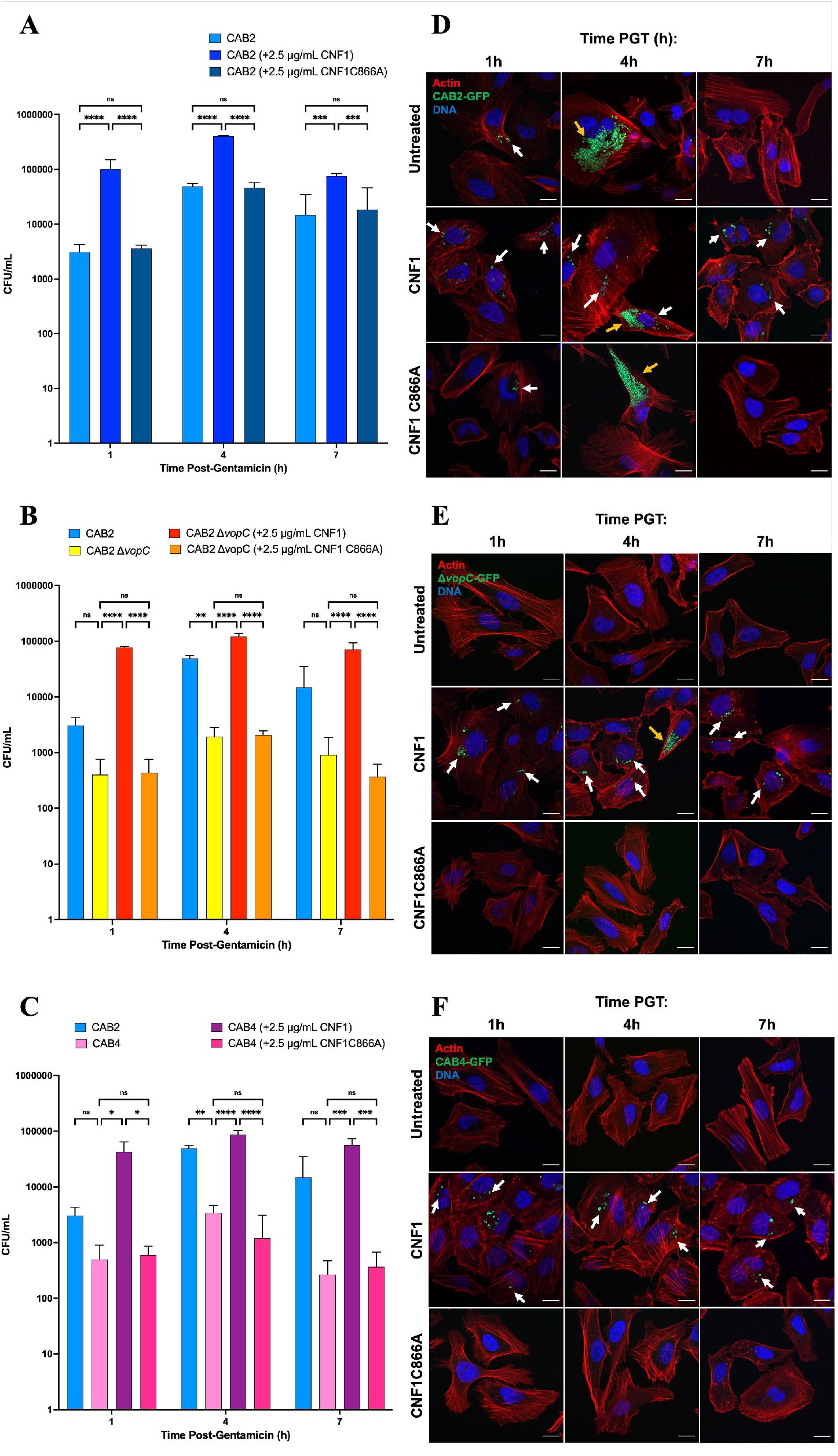
CNF1-mediated invasion is independent of VopC and the T3SS2. **(A-C)** Gentamicin protection assay comparing intracellular **(A)** CAB2, **(B)** CAB2Δ*vopC*, or **(C)** CAB4 at 1, 4, and 7 hour post-gentamicin application, which proceeded infection and mock, 2.5μg/mL CNF1, or 2.5μg/mL CNF1 C866A application by 2 hours. All infections were conducted at an MOI = 10 Error bars represent standard deviation of three technical replicates. Statistical significance measured using a two-way ANOVA with multiple comparisons test (ns, not significant; *, P < 0.05; **, P < 0.005; ***, P < 0.0005; ****, P < 0.00005). Gentamicin protection assay quantifications are each representative of three biological replicates **(D-F)**. Representative confocal micrographs of HeLa cells infected with **(D)** CAB2-GFP, (**E)** CAB2Δ*vopC*-GFP, or **(F)** CAB4-GFP at an MOI = 10 and untreated (top) or treated with 2.5μg/mL CNF1 (middle) or 2.5μg/mL CNF1 C866A (bottom) for 2 hours before application of gentamicin. All HeLa cells were stained for F-actin using rhodamine-phalloidin (red) and Hoechst DNA stain (blue). White arrows indicate endosomal internalized CAB2-GFP. Yellow arrows indicate cytoplasmic internalized CAB2-GFP. Scale bar, 20μm.

At 4 hours PGT, we observed an increase in intracellular bacteria for all treatment conditions, consistent with CAB2’s canonical escape from the endosome and cytoplasmic replication at this time point (Fig. 3A, D, Table S1) (31). One notable difference observed between the untreated and CNF1-treated infection conditions, however, was the abundance of GFP positive puncta within the HeLa cells after CNF1 treatment. The bacteria clustered in these puncta strongly resembled those visible at 1 hour PGT, when CAB2 would normally be endosomal, suggesting that CNF1-treated cells had aborted their infection at endosomal stage after the initial invasion step (Fig. 3D). This hypothesis was corroborated when we observed that CNF1 C866A treatment did not culminate in higher intracellular CFUs than the untreated control CAB2 infection (Fig. 3A, D). We observed that almost all the bacteria infected with CAB2 or CAB2 in the presence of CNF1 C866A transitioned to more dispersed cytoplasmic growth (Fig. 3A, D). Finally, the continued presence of GFP positive bacteria in puncta, in conjunction with the persistently elevated CFU count in the CNF1-treated conditions at 7 hours PGT, suggested that CAB2 infection was arrested at endosomal stage in the presence of CNF1 (Fig. 3A, D).

### CNF1-mediated invasion is not contingent on VopC or the T3SS2

Though most CAB2 internalized after CNF1 treatment remained endosome-bound, a relatively small subset of invaded bacteria did appear capable of cytosolic proliferation and completing their infection cycle (Fig. 3D). In a few invaded cells treated with CNF1 we observed the same “dispersed” bacterial distributions reminiscent of cytosolic replication in the canonical *V. parahaemolyticus* life cycle at 4 hours PGT (Fig. 3D). Furthermore, bacteria in the dispersed growth morphology were largely absent in the CNF1-treated condition by 7 hours PGT, while those in the punctate growth morphology remained, suggesting bacteria capable of egressing from the endosome could egress out of the cell also (Fig. 3D). Since VopC is canonically responsible for driving *V. parahaemolyticus* invasion, and *V. parahaemolyticus* canonically escapes the endosome by 7-hours PGT, we sought to determine whether CNF1-mediated invasion was VopC-dependent. We hoped also to address whether the small population of bacteria apparently able to complete their infection cycle after CNF1 treatment could do so independently of VopC.

Thus, we conducted another gentamicin protection assay, this time including alongside CAB2 both the CAB2Δ*vopC* mutant strain, lacking the VopC effector required for invasion, and the CAB4 strain, lacking hemolysins and the transcriptional regulators for T3SS1 and T3SS2 expression (4). At 1, 4, and 7 hours PGT, CNF1 treatment significantly increased the quantity of intracellular bacteria for all three strains of bacteria relative to the untreated and the CNF1 C866A-treated controls, both of which exhibited similar quantities of intracellular CFUs (Fig. 3A-C). We also once again examined HeLa cells infected with CAB2-GFP, CAB2Δ*vopC*-GFP, and CAB4-GFP via confocal microscopy. The infection profiles of CAB2 and CAB2Δ*vopC*-GFP in the presence of CNF1 bore significant similarities. Both exhibited many more GFP-positive bacteria forming puncta within HeLa cells at 1 hour PGT than were observed in the untreated and CNF1 C866A-treated controls. Moreover, this increased presence of punctate bacteria persisted at 4 and 7 hours PGT after CNF1 treatment. At 4h PGT, however, a small subset of CAB2Δ*vopC*-GFP were observed growing in dispersed clusters reminiscent of the cytosolic growth observed in CAB2-GFP at the same time point. That these dispersed clusters represented cytosolic growth was evidenced also by their disappearance by 7 hours PGT, indicating egress from HeLa cells (Fig. 3D-E).

In contrast with CAB2-GFP and CAB2Δ*vopC*-GFP, CAB4-GFP in the presence of CNF1 appeared almost exclusively as puncta at all three time points (Fig. 4F). In the untreated and CNF1 C866A-treated control conditions for CAB2Δ*vopC* and CAB4, no infected cells were observed under confocal microscopy, and significantly fewer intracellular CFUs were observed in gentamicin protection assays than were counted in the presence of CNF1. (Fig. 3B-C, E-F).

**Fig 4.**
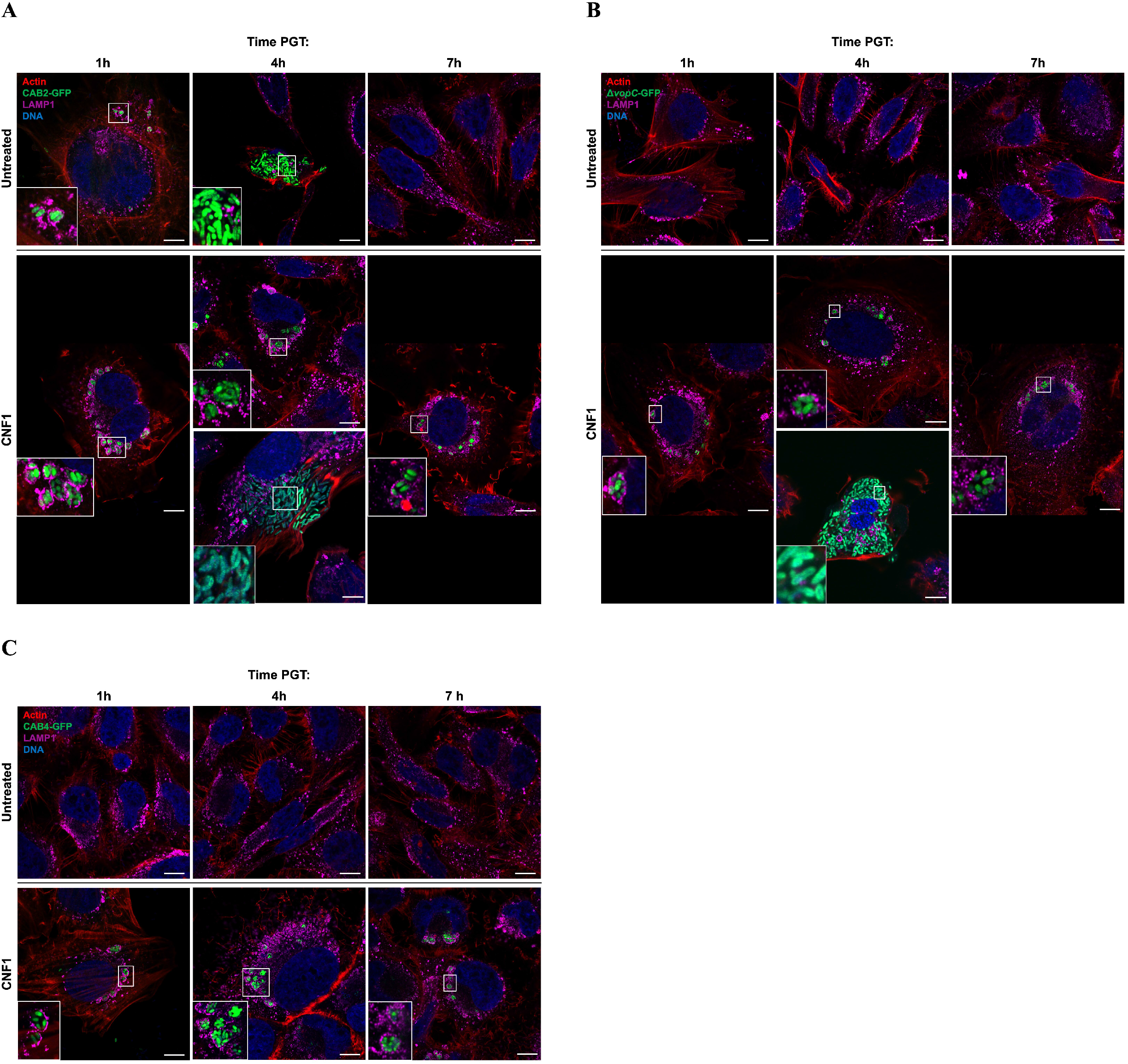
*V. parahaemolyticus* puncta are endosome-bound, while escaped *V. parahaemolyticus* grow in the cytoplasm. **(A-C)** Representative confocal micrographs of HeLa cells infected for 2 hours with **(A)** CAB2-GFP, **(B)** CAB2Δ*vopC*-GFP, or **(C)** CAB4-GFP at an MOI = 10 and untreated (top) or treated with 2.5μg/mL CNF1 (bottom) for 2 hours before application of gentamicin. **(A &B)** Images of both endosomal and cytoplasmic bacteria in CNF1-treated cells at 4 hour PGT are shown. White boxes demarcate magnified area in the bottom left of the corresponding image. All HeLa cells were stained for F-actin using Alexafluor680-phalloidin (red), for late endosomes with lamp-1 antibodies (purple), and DNA with Hoechst (blue). Scale bar, 10μm.

### *V. parahaemolyticus* internalized after CNF1 treatment remain largely confined to the endosome

After identifying two primary intracellular bacterial morphologies, punctate and dispersed, associated with CNF1 treatment, we tested whether these morphologies truly represented endosomal and cytoplasmic replication inside host cells. Because *V. parahaemolyticus*’s intracellular life cycle begins within the endosome, and escape into the cytoplasm is preceded by endosome maturation, we immunofluorescently stained for lysosomal-associated membrane protein -1 (Lamp-1), a late endosome marker, in infected HeLa cells for confocal microscopy analysis (31). CAB2-GFP, CAB2Δ*vopC*-GFP, and CAB4-GFP in the punctate morphology were surrounded by LAMP-1-laden endosomes after CNF1 treatment at 1, 4, and 7 hours PGT (Fig. 4, insets). This LAMP-1 staining profile was identical to that observed in CAB2-GFP infections without CNF1 at 1 hour PGT, when the endosomes surrounding bacteria are coated in LAMP-1 (Fig. 4A) (31). The dispersed morphology of CAB2-GFP at 4 hours PGT in the absence of CNF1 lacks LAMP-1 staining around the bacteria, indicating a cytosolic locus of replication (Fig. 4A). This in turn suggests CAB2-GFP and CAB2Δ*vopC*-GFP in the dispersed morphology are also replicating cytosolically, consistent with the canonical *V. parahaemolyticus* T3SS2-mediated infection cycle, as they too lack LAMP-1 staining around the periphery of the bacterial growth (31) (Fig. 4A-B).

Having qualitatively characterized the dynamics of CNF1 invasion by linking the CNF1-associated punctate morphology with endosomal localization, we adopted a more quantitative approach to investigate CNF1-mediated invasion. Although gentamicin protection assays in Figures 3 and 4 demonstrated the capacity of CNF1 to increase invasion, we still had not determined whether this increase was attributable to more HeLa cells infected, more invasion events per cell, or a combination of the two. By extension, we also hoped to determine whether, after CNF1 treatment, CAB4 remained confined to the endosome after invasion more frequently than CAB2 and CAB2Δ*vopC*.

Using confocal microscopy, we first approximated the percentage of HeLa cells infected by CAB2-GFP, CAB2Δ*vopC*-GFP, and CAB4-GFP with and without CNF1 or CNF1 C866A. We confirmed across multiple trials that CAB4 and CAB2Δ*vopC* were unable to invade at all without CNF1 present (Fig. 3B-C, E-F). However, HeLa cells were invaded by CAB2Δ*vopC* and CAB4 at 1 hour PGT in the CNF1-treated condition (Fig. 3B-C, E-F). Though somewhat less pronounced, CAB2 also invaded more cells on average in the presence of CNF1 than in its absence, as was the case for CAB2Δ*vopC* and CAB4 at 1, 4, and 7 hours PGT (Fig. 3A, D). Moreover, we determined that CAB4 invaded slightly fewer cells than did CAB2 or CAB2Δ*vopC*, which exhibited roughly equal invasion frequency at 1-hour PGT, though this finding did not translate to significantly fewer intracellular CFUs recovered in the gentamicin protection assays shown in Figures 3A-C (statistical comparison not shown).

Comparisons between strains are shown at 4 hours PGT, when endosomal escape normally peaks in the canonical *V. parahaemolyticus* life cycle (31). As expected, 100% of intracellular bacteria for all strains and treatment conditions were endosomal at 1 hour PGT. The number of endosome-bound bacteria dropped between 1 and 4 hours PGT for CAB2 in all treatment conditions, though a significantly higher fraction of intracellular CAB2 remained endosome-bound in the CNF1-treated condition at 4 hours PGT than in the control conditions (Fig. 5). After CNF1 treatment, CAB2Δ*vopC* also exhibited a small drop in the fraction of endosome-bound bacteria between 1 and 4 hours PGT comparable to that of CNF1-treated CAB2, with most of intracellular bacteria remaining endosome-bound at 4 hours PGT, and a small minority escaping to the cytosol (Fig. 5). CAB4 appeared largely unable to escape the endosome at all by 4 hours PGT (Fig. 5). At 7 hours PGT, no intracellular CAB2 was observed in the control conditions, as the bacteria had by that time completed their canonical infection cycle and escaped the host (31). In contrast, CNF1-treated CAB2, CAB2 Δ*vopC*, and CAB4 all appeared exclusively endosome-bound by 7 hours PGT (data not shown). The absence of cytoplasmic CAB2 and CAB2Δ*vopC* at 7 hours PGT suggested that the cytoplasmic bacteria observed after CNF1 treatment at 4 hours PGT could successfully complete their infection cycle, and thus could escape the host by the 7 hour time point. The absence of cytoplasmic bacteria of any strain, alongside the continued presence of endosomal bacteria, at 7 hours PGT in the CNF1 treatment condition, also suggested that bacteria unable to escape the endosome by, roughly, 4 hours PGT would be unable to escape the endosome at all.

**Fig 5.**
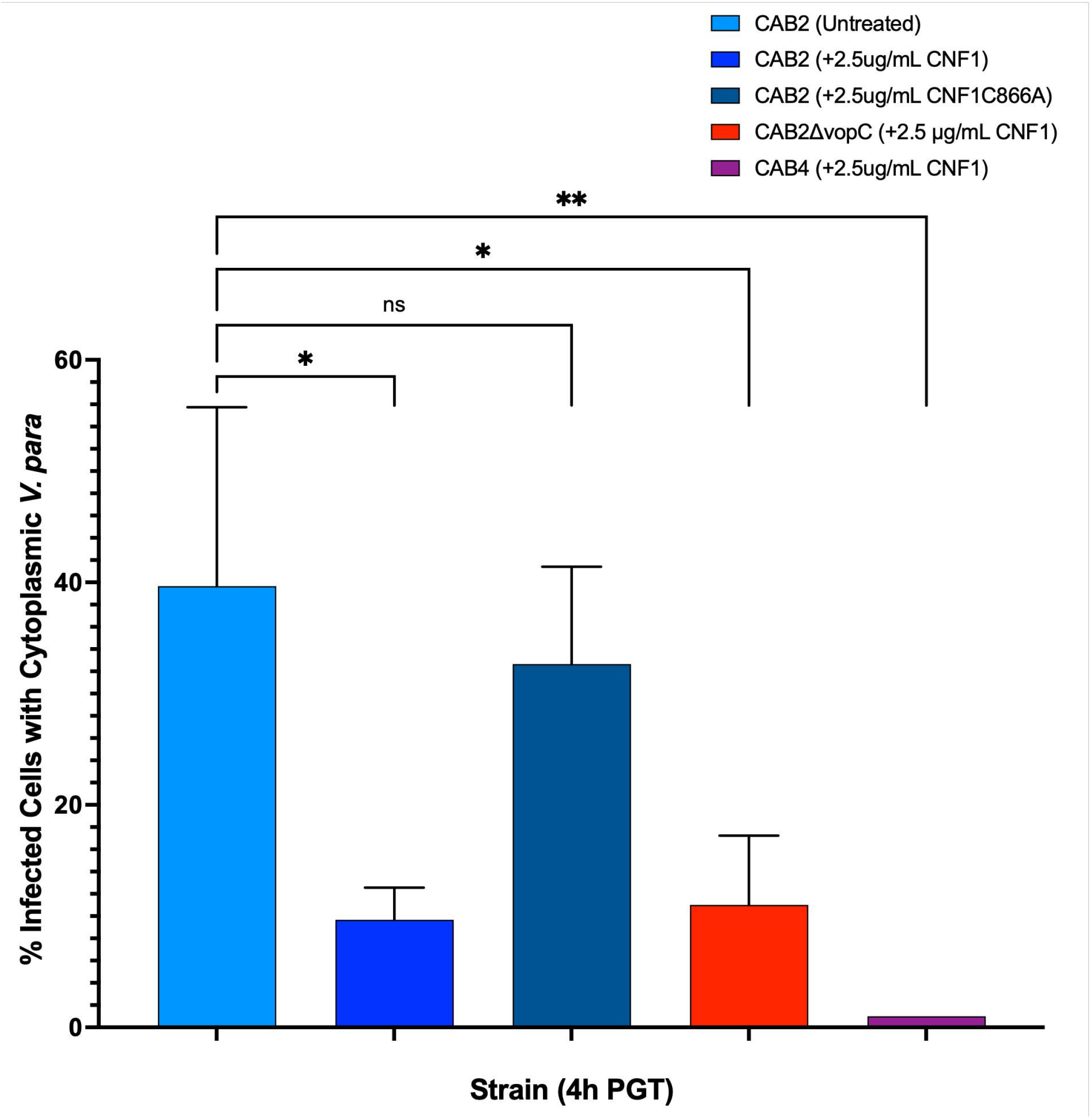
Comparison of the percentage of infected HeLa cells containing cytoplasmic *V. parahaemolyticus*, based on confocal microscopy. HeLa cells infected for 2 hours with CAB2-GFP, CAB2Δ*vopC*-GFP, or CAB4-GFP at an MOI = 10 and untreated or treated with 2.5μg/mL CNF1 or 2.5μg/mL CNF1 C866A for 2 hours before application of gentamicin. Each percentage was calculated for each strain by averaging the percentage of infected HeLa cells with cytoplasmic *V. parahaemolyticus* out of the total infected cells on a cover slip. Percentages from three separate infection experiments were averaged and displayed here. Statistical significance measured using one-way ANOVA with multiple comparisons test (ns, not significant; *, P < 0.05; **, P < 0.005).

**Fig 6.**
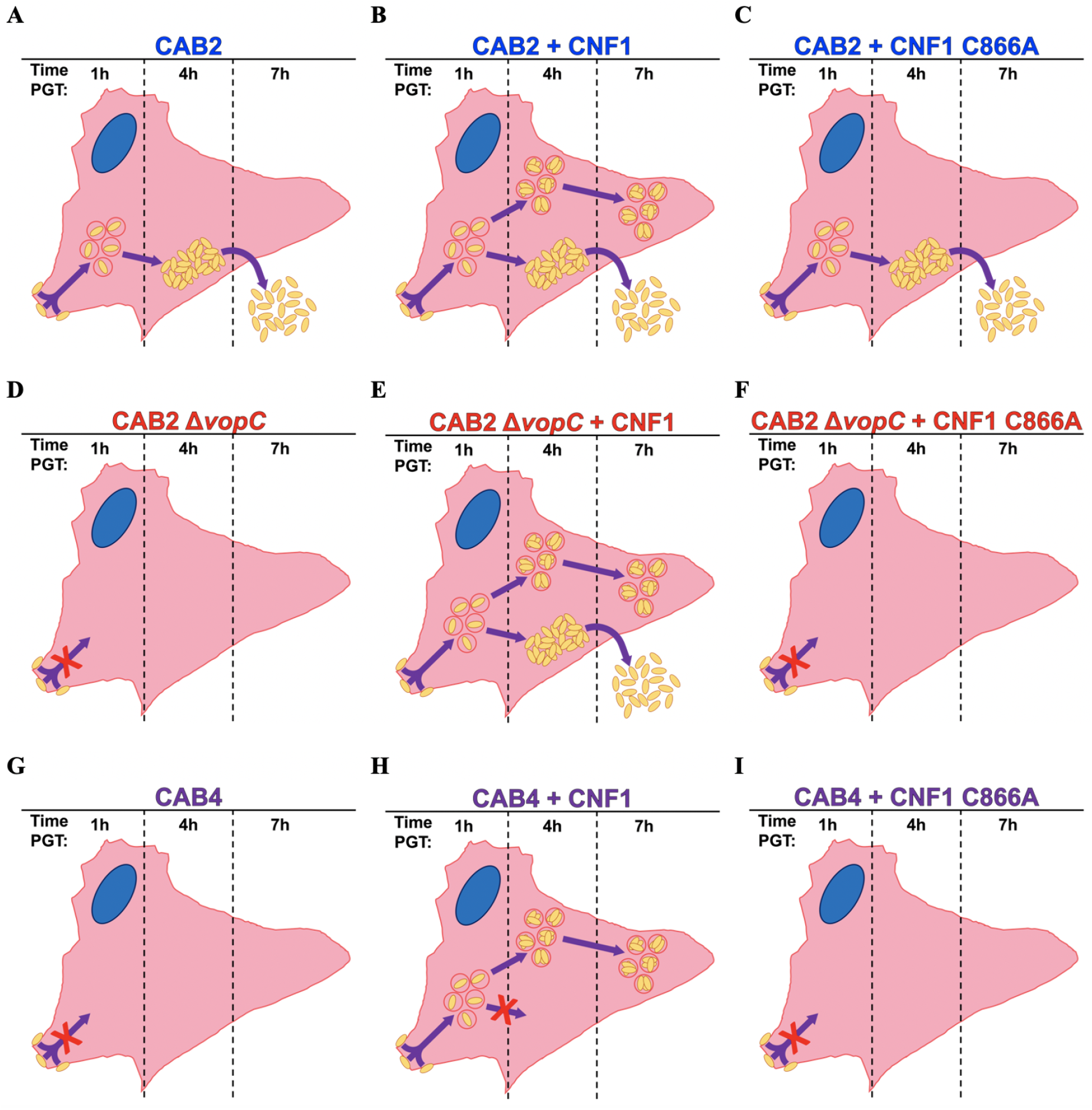
Model of infection phenotypes for *V. parahaemolyticus* strains with and without CNF1. **(A-C)** CAB2 infection progression. CAB2 adheres to the host and invades in a VopC- or CNF1-mediated mechanism by 1 hour PGT. Bacteria escape from the endosome into the cytoplasm by 4 hours PGT, and ultimately escape from and kill the host-cell by 7 hours PGT. In the presence of CNF1, a majority of infected cells exhibit no endosomal escape at 4 hours PGT, and these bacteria remain confined to the endosome even by 7 hours PGT. **(D-F)** CAB2Δ*vopC* infection progression. Due to the absence of VopC, CAB2Δ*vopC* is unable to invade the host without biochemical supplementation, but is able to invade the host in the presence of CNF1. Bacteria escape from the endosome into the cytoplasm by 4 hours PGT, and ultimately escape from and kill the host-cell by 7 hours PGT. In the presence of CNF1, a majority of infected cells exhibit no endosomal escape at 4 hours PGT, and these bacteria remain confined to the endosome even by 7 hours PGT. **(G-I)** CAB4 infection progression. Due to the absence of the T3SS2, CAB4 is unable to invade the host without biochemical supplementation, but is able to invade the host in the presence of CNF1. No bacteria were observed escaping from the endosome by 4 hours PGT after CNF1 application while internalized bacteria persisted within endosomes 4 hours PGT and 7 hours PGT.

## Discussion

*V. parahaemolyticus* exhibits low baseline levels of invasion *in vitro*, but is nonetheless a leading cause of foodborne acute gastroenteritis with an ever-expanding ecological range as coastal waters warm (5, 35, 36). Like many Gram-negative bacteria, *V. parahaemolyticus* infection is predicated upon the deamidation of host Rho GTPases; however, despite the similarities between mechanisms of entry across bacterial species, relatively little research has focused on cross-species compatibility of these enzymes (4, 17, 34). The secreted toxin CNF1 and T3SS-translocated effector VopC are homologous Rho GTPase deamidases that mediate the invasion of the UPEC and *V. parahaemolyticus*, respectively (Fig. 1, S1) (4, 5). Because of this homology, we hypothesized CNF1 could markedly increase invasion levels of *V. parahaemolyticus* above those mediated by VopC (4). To test our hypothesis, we first investigated whether CNF1 could improve *V. parahaemolyticus* invasion in a tissue culture model. We found that CNF1 increased the frequency of *V. parahaemolyticus* invasion, but that VopC, despite its inefficiency, facilitates the full course of *V. parahaemolyticus* infection in ways that CNF1 does not. In fact, although CNF1 drastically increased *V. parahaemolyticus* invasion frequency, most endosomal *V. parahaemolyticus* aborted their infection process after CNF1-mediated invasion, and never escaped from the endosome into the host cytoplasm (Fig. 3B, 7). We also determined that CNF1 drives *V. parahaemolyticus* invasion in the absence of both VopC and the T3SS2 altogether, and confirmed CAB2, CAB2Δ*vopC*, and CAB4 all remained largely unable to progress past the invasion stage of infection after CNF1 treatment (Fig. 3, 5). A closer examination of these infections revealed that a small-but-significant fraction of CAB2 and CAB2Δ*vopC* did, in fact, complete their infection cycle after CNF1 treatment, while CAB4 remained almost entirely endosome-bound for the entirety of the infection (Figs. 4 and 5). In addition to demonstrating the capacity of CNF1 to promote invasion in a VopC- and T3SS-independent manner, these experiments indicated that endosomal escape was VopC-independent. The absence of cytoplasmic CAB4-GFP during these infections, along with the presence of cytoplasmic CAB2 and CAB2Δ*vopC*, point to the necessity of the VtrABC signaling cascade, the regulatory apparatus of the T3SS2 and associated virulence factors disrupted in CAB4, but intact in CAB2 and CAB2Δ*vopC*, in mediating endosomal escape, as well as T3SS2 expression (4, 37).

In examining the infection dynamics of bacteria internalized after CNF1 treatment, we concluded, based on the inability of CNF1 C866A to promote invasion in the absence of VopC or to increase invasion above untreated levels, that the CNF1-dependent increase in the invasion of HeLa cells by CAB2, CAB2Δ*vopC*, and CAB4 was attributable to the catalytic activity of the CNF1 (Fig. 3). We could also deduce from this the substitution of cysteine 866 in the CNF1 active site with an alanine was sufficient to disrupt the enzyme’s ability to promote bacterial invasion (Fig. 3). Moreover, since untreated and CNF1 C866A-treated CAB2 had egressed from nearly all cells by 7 hours PGT, and because the presence of gentamicin in the medium prevented escaped CAB2 from re-infecting new HeLa cells, we determined that bacteria confined to the endosome by 4 hours PGT after CNF1-mediated infection remained permanently endosome-bound unless killed by fusion with the lysosome (Fig. 3D). This hypothesis held true for both CAB2Δ*vopC* and CAB4 after treatment with CNF1 as well, since almost all CAB4 remained endosome-bound at all time points, and while cytoplasmic CAB2Δ*vopC* were observed at 4 hours PGT alongside many endosomal bacteria, all intracellular CAB2Δ*vopC* observable by 7 hours PGT were endosomal (Fig. 3E, F).

Our quantitative analyses counted significantly more intracellular CAB2 and CAB2Δ*vopC* at 1 hour PGT than we did CAB4 at the same time point (Fig. 3A-C). Because CAB2 and CAB2Δ*vopC* at 1 hour PGT after CNF1 treatment did not differ significantly from one another, we could rule out the possibility of native VopC contributing significantly to the aforementioned difference in invasion levels. With CNF1 established as the primary diver of invasion, we speculate that the differences observed in intracellular bacterial counts at 1-hour PGT were rooted not in any difference in the internalization efficiency across strains, but the number of bacteria available for internalization in the first place. It is likely that CAB4’s reduced intracellular bacterial counts after CNF1 treatment rest in some defect in adhesion to the host. Corroborating this notion is the established role of T3SS translocon pores in adhesion; thus, because CAB4 cannot produce any T3SS components (to say nothing of other potential adhesion factors regulated by the VtrABC signaling cascade), and adhesion is a well-established prerequisite for invasion, CAB4 may, of the strains tested, uniquely lack a full complement of adhesion machinery required for efficient invasion (38, 39).

More pertinent to our original research question is our observation that the fates of *V. parahaemolyticus* internalized via CNF1 differ substantially from those which invade in a VopC-dependent manner. At the most basic level, the issue may rest simply in the capacity of internalized bacteria to translocate effectors at all. Ordinarily, *V. parahaemolyticus* are able to invade only after the T3SS2 has engaged with the host plasma membrane and translocated VopC into the host cytosol (4). *V. parahaemolyticus* is able to adhere to host cells independently of the T3SS2, however, suggesting that the addition of CNF1 may promote invasion of bacteria that have adhered to the host plasma membrane, but have not engaged the T3SS2 and translocated effectors (40–42). Since endosomal escape appears to hinge on VtrABC signaling, and plausibly on the T3SS2 itself, many of the bacteria internalized by CNF1 may have been so prematurely, without the T3SS2 engaged, and are thus unable to deliver the virulence factors necessary to promote endosomal escape into the host cytosol.

Another potential culprit behind the inability of *V. parahaemolyticus* internalized by CNF1 to escape the endosome is the difference in specificity of CNF1 and VopC. Both VopC and CNF1 target Rac1 and CDC42, but only CNF1 deamidates RhoA (Fig. 1). Although CDC42, Rac1, and RhoA are highly conserved and all regulate the actin cytoskeleton in a general sense, the differences between them are not negligible. Active Rac1 and CDC42 promote filament stabilization by binding and activating the kinase PAK1, which phosphorylates and activates LIM-K kinases responsible for deactivating the actin depolymerizing enzyme cofilin through phosphorylation (43, 44). These proteins in their GTP-bound state also promote the nucleation and elongation of actin branches, rather than *de novo* actin filaments, respectively by interacting with WASP and WAVE proteins that bind and recruit the ARP2/3 complex (45–47). Active RhoA, by contrast, promotes actin polymerization and filament elongation through the upregulation of formins such Daam and mDia, which nucleate and polymerize monomeric actin into new filaments (48, 49). RhoA is also able to stabilize these filaments through the phosphorylation and activation of LIM-K, as are Rac1 and CDC42; however, RhoA does so by engaging a different signaling pathway than Rac1 and CDC42, as it binds and activates RhoA-associated kinases, ROCKs, to phosphorylate and activate LIM-K, rather than signaling through PAK (43, 50).

As many *V. parahaemolyticus* virulence factors target the actin cytoskeleton, it stands to reason that even slight differences in substrate specificity between VopC and CNF1 could culminate in significantly different outcomes for the bacteria during infection. VopC evolved to function against the backdrop of a very specific battery of effectors targeting the actin cytoskeleton. For example, the *V. parahaemolyticus* T3SS2 effector VopL drives the polymerization of nonfunctional actin fibers through its three WH2 domains, while the catalytic activity of the T3SS2 effector VopV binds and bundles actin filaments during infection (12, 51). The activities of both effectors could conflict with those of an activated RhoA. The nonfunctional VopL-mediated actin fibers are intended to sequester actin, and thus may not be sufficient for doing so when RhoA is driving actin filament polymerization at the same time. Likewise, VopV’s actin bundling activity is not dissimilar to that of RhoA-regulated myosin, and thus the upregulation of myosin-mediated actin bundling may compete or conflict with VopV’s intended targets (12, 52, 53).

Another interesting observation is the presence of a DUF4765 domain, a putative ADP-ribsoyltransferase domain, in D4 of CNF1 (Fig S1A). The role of this domain and its contributions to the infection process of UPEC are not known, but its fusion to the deamidase catalytic domain of CNF1 ensures that both enzymes are proximal to one another throughout the UPEC infection process (20). *V. parahaemolyticus* also translocates an ADP-ribosyltransferase into the host as an effector the T3SS2, VopT (10). Thus while both the CNF1 and VopC deamidases are likely functioning in the presence of an ADP-ribosyltransferase, CNF1 is (albeit putatively) fused to this enzyme, wherease VopC is simple translocated alongside it. While further research is necessary to elucidate the target and catalytic activity of the CNF1 D4 domain, the question of whether or how colocalization of an ADP-ribosyltransferase with a deamidase affects the fate of intracellular bacteria is an interesting and open one (5, 10, 20).

The differences between the intracellular environment after VopC-mediated invasion and CNF1-mediated invasion may be especially stark, since VopC should, in theory, repress RhoA activity, while CNF1 constitutively activates RhoA. Rac1 has been shown to repress RhoA signaling, and not only can CDC42 promote Rac1 activation, but both Rac1 and CDC42 have been shown specifically to block the phosphorylation and activation of myosin through PAK signaling as well (53, 54). The picture is complicated by the T3SS2 effector VopO, which acts as a GEF of RhoA and has been linked to maintaining the efficiency of *V. parahaemolyticus* invasion; however, VopO may act as a precise counterbalance to the predicted downstream effects of VopC. Since Rac1-mediated repression of RhoA is thought to occur through downregulation of GEFs, *V. parahaemolyticus* may leave only a relatively few translocated VopO molecules to maintain RhoA activity, possibly (though speculatively) at a level well below what could be achieved by constitutive activation through deamidation (13, 53). A hypothesis concerning the interplay of these virulence factors might posit that VopC strongly depresses RhoA activity, while VopO serves as a relatively weak counterbalance to VopC against the backdrop of an actin pool depleted by VopL and further modulated by VopV. Within such a model, CNF1 might would function like a thumb on the scale of the delicate balance of *V. parahaemolyticus*’s modulation of actin dynamics, leading to the arrest of *V. parahaemolyticus* infection at the endosomal stage.

The reason underlying the incompatibility of CNF1 with the *V. para* infection cycle is not unprecedented in the bacterial world, or even within *V. para*. The *V. para* T3SS2 effector VopA, for example, is a serine, threonine, and lysine acetylase with significant homology to the *Yersinia* T3SS effector YopJ (8, 51). The key differences between these effectors rests not in their catalytic activities, but in their selective targeting of substrates, analogous the differences in small Rho GTPase targeting between CNF1 and VopC. YopJ possesses the broader range of targets of the two, acetylating and blocking regulatory kinases that regulate JNK, ERK, p38, and NFkB signaling (8, 51). By broadly blocking MAPK signaling, YopJ shuts down the host innate immune response by preventing cytokine induction (51, 52). By specifically crippling the NFkB pathway, however, YopJ also impairs host anti-apoptotic signaling, promoting cell death (51, 52). *Yersinia* is an extracellular pathogen, and thus stands to benefit from the nutrients released by host apoptosis, just as its survival is enhanced by impairing host innate immune signaling (52). *V. para*, by contrast, replicates intracellularly, and while it, like *Yersinia*, must evade the host innate immune response to survive, it cannot compromise host survival without also compromising its intracellular replicative niche (27, 52). It should come as little surprise, then, that VopA has evolved to target MAPK signaling less broadly than YopJ, and has been shown to acetylate the regulatory kinases of the JNK, ERK, and p38 signaling pathways, but not those regulating NFkB (8, 53). Thus while *V. para* blocks the MAPK signaling pathways promoting cytokine induction and apoptosis signaling, likely allowing the intracellular *V. para* to evade host innate immunity, it leaves the NfKB signaling pathway untouched, avoiding apoptosis induction and the premature destruction of its intracellular replicative niche (8). While speculative, it is nonetheless plausible that the target specificity of VopA relative to its close homolog YopJ parallels that of VopC relative to CNF1, and the relationship between VopA and YopJ highlights how differences in bacterial lifestyle can culminate in divergent evolutionary pressures on virulence factors, which consequently affect the host in radically different ways despite retaining significant homology to one another.

Ultimately, our data demonstrate both the efficacy and the limitations of leveraging conserved invasion mechanisms to increase *V. parahaemolyticus* invasion *in vitro*. Even as the percentage of invaded HeLa cells increased significantly after the application of CNF1, the percentage of invaded cell that exhibited aborted *V. parahaemolyticus* infections also increased, demonstrating that for *V. parahaemolyticus*, CFN1’s utility as a tool promoting infection efficiency is severely limited. Therefore, the soluble factor CNF1 secreted by UPEC can mediate invasion for neighboring pathogens, but this is unlikely to result in successful virulence. These experiments clearly illustrate the incredible precision that individual bacteria have evolved in their infection processes, such that even highly conserved mechanisms of invasion and infection are not always compatible across species. What remains are many, now-refined questions about the interplay of regulation and virulence factors responsible for each step of infection that future experiments will need to address.

## Materials and Methods

### Strains and plasmids

The *V. parahaemolyticus* CAB2 strain was derived from POR1 (clinical isolate RIMD2210633 lacking TDH toxins) and contains an additional deletion for the gene encoding ExsA, the transcription factor responsible for activating the T3SS1 (4). CAB2Δ*vopC* was derived from CAB2, containing an additional deletion in the coding sequence for the T3SS2 effector VopC (4). CAB4 was derived from POR1, containing two additional deletions for the gene encoding ExsA, as well as VtrA, the transcription factors responsible for activating the T3SS1 and T3SS2 respectively (4). CAB2-GFP, CAB2Δ*vopC*-GFP, and CAB4-GFP were generated via a standard triparental mating that transformed each *V. parahaemolyticus* strain with the pMW-GFP vector (58). All strains were grown aerobically in Luria-Bertani (LB) medium, supplemented with NaCl to a final concentration of 3% (w/v) (marine LB, or MLB) at 30°C. Strains expressing GFP were grown under identical conditions, with the addition of 50 μg/mL spectinomycin.

### Expression and purification of CNF1

CNF1 was expressed from *E*.*coli* BL21 transformed with pET28a containing a copy of *cnf1* as sequenced by Falbo, *et al*. with an N-terminal 6xHis tag (59). The vector was synthesized by Twist Bioscience. BL21 containing the CNF1 expression vector were grown to exponential growth phase in 2xyT media aerobically at 37°C, and expression of CNF1 was induced overnight in 0.4 mM isopropyl-β-D-thiogalactopyranoside aerobically at 20°C. Induced bacteria were pelleted and lysed via sonication, and clarified lysates were incubated with Qiagen NiNTA resin (30210) nutating at 4°C. Protein-bound resin was transferred to a column, which was subsequently washed with 20 mM imidazole in 50mM Tris-HCl pH8.0, 150mM NaCl, and the toxin was eluted with 250 mM imidazole in 50mM Tris-HCl pH8.0, 150mM NaCl. The toxin was subsequently buffer exchanged via gel filtration chromatography (Superdex 200 HiLoad 16/600 **GE28-9893-35**) in 50mM Tris-HCl pH8.0, 150mM NaCl, and concentrated to 5mg/mL in an Amicon 100kDa spin filter at 4°C. The toxin was aliquoted, flash frozen in liquid nitrogen, and kept at -80°C for storage. CNF1 C866A was generated via PCR mutagenesis using primers designed with Agilent Quickchange Primer design, pET28a with *cnf1* as a template, and Thermo Phusion polymerase.

### Gentamicin protection assays

HeLa cells were plated in triplicate in a 96-well tissue culture plate at 7*10^4^ cells/ml per well and grown for 16-18h. Bacteria were added to triplicate wells of HeLa cell monolayers for infection in media containing no additives, 2.5µg/mL CNF1, or 2.5µg/mL CNF1 C866A. All infections were carried out at a MOI of 10, and bacteria were induced for 1.5 h in MLB media supplemented with 0.5% w/v bile salts prior to infection (Sigma Aldrich B3883). Infections were synchronized by centrifugation at 1000xg for 5 minutes after addition of bacteria to wells, and allowed to infect HeLa cells for 2 hours. Gentamicin was added at 100µg/mL to each well after 2 h of infection to kill extracellular bacteria. At each indicated time point, monolayers of HeLa cells were washed with PBS and cells were lysed by incubation with PBS containing 0.5% Triton X-100 for 10 min at room temperature with agitation. Serial dilutions of lysates were plated on MMM (minimal marine medium) plates and incubated at 30 °C overnight for subsequent CFU enumeration.

### Infection assays for confocal imaging

HeLa cells were seeded onto 6-well plates containing sterile coverslips at a density of 7*10^4^ cell/mL. Following infections with *V. parahaemolyticus* strains at an MOI of 10 and addition of CNF1/ CNF1 C866A as detailed above, cells were washed with PBS and fixed in 3.2% (v/v) paraformaldehyde for 10 min at room temperature. Fixed cells were washed in PBS and permeabilized with 0.1% Triton X-100 for 10 min at room temperature. Nuclei and actin cytoskeleton were stained with Hoechst 33342 (Sigma) and rhodamine-phalloidin (Molecular Probes), respectively, for infection analyses and quantification. For evaluating endosomal localization of bacteria, nuclei, actin cytoskeleton, and Lamp-1 were stained with Hoechst 33342 (Sigma), Alexafluor680-phalloidin (Molecular Probes), and Mouse anti-Lamp1 (Abcam Ab25630) respectively as described previously (31). Images were collected using a Zeiss LSM 710 confocal microscope.

### Quantification of confocal images

The assessment of whether bacteria were endosomal or cytoplasmic was predicated on whether the bacteria were growing in the punctate or dispersed morphology as described above. The fractions of infected cells containing exclusively endosomal bacteria out of the total number of infected cells on each cover slip were collected for each bacterial strain, time point, and treatment (either with nothing, CNF1, or CNF1 C866A), and the fractions for each condition averaged across three experiments. All quantifications were collected blind to the identity of each sample.

### Growth curve

Strains were grown overnight shaking at 30°C in MLB supplemented with 50µg/mL spectinomycin. For comparison of growth rates, overnights were used to inoculate 50mL MLB supplemented with 50µg/mL spectinomycin at a starting OD_600_ = 0.02 in triplicate. Cultures were incubated at 30°C shaking, with OD_600_ measurements collected hourly for 8 hours, and once more after 32 hours. Each data point represents an average of three technical replicates.

### Statistical analysis

Unless otherwise stated, all data are presented as mean ± standard deviation of three or more independent experiments. All experiments discussed in this paper were conducted in triplicate. A two-way ANOVA with Tukey’s multiple comparisons test was conducted to evaluate statistical significance between all variables. A P value of < 0.05 was deemed significant.

## Acknowledgements

We thank the Orth lab members for discussions and editing. This work was funded by the Welch Foundation grant I-1561 (K.O.), Once Upon a Time…Foundation (K.O.), and National Institutes of Health Grants 5T32AI007520-20 (A.L.), R01 GM115188 (K.O.). K.O. is a W.W. Caruth, Jr. Biomedical Scholar with an Earl A. Forsythe Chair in Biomedical Science.

## Table Legends

**Table S1** Expanded list of gentamicin protection assay statistical comparisons from Figure 3. Briefly, gentamicin protection assay comparing intracellular CAB2, CAB2Δ*vopC*, or CAB4 at 1, 4, and 7 hour post-gentamicin application, which proceeded infection and mock, 2.5μg/mL CNF1, or 2.5μg/mL CNF1 C866A application by 2 hours. All infections were conducted at an MOI = 10 Statistical significance measured using a two-way ANOVA with multiple comparisons test (ns, not significant; *, P < 0.05; **, P < 0.005; ***, P < 0.0005; ****, P < 0.00005)

## Supplemental Figure Legends

**Fig S1** Comparison of *V. parahaemolyticus* VopC and UPEC CNF1. **(A)** Schematic of VopC (top) and CNF1 (bottom) domain boundaries. CBD: chaperone binding domain. Numbers above each schematic correspond to amino acid position. **(B)** Local sequence alignment of VopC and CNF1 catalytic domains. Sequences are labeled according to species to the left of the alignment, and by sequence number to the left and right. Conserved residues are boxed, with identical amino acids highlighted in red and similar residues unhighlighted. Asterisks indicate conserved catalytic residues. Alignment generated using ESPript3.0 (61) **(C)** Structural prediction of the predicted VopC catalytic domain model side-by-side with the CNF1 catalytic domain crystal structure (PDB 1hq0). Conserved catalytic cysteine and histidine are identified in both proteins. **(D)** Comparison of the predicted VopC catalytic domain structure (cyan) and CNF1 catalytic domain structure (orange). Conserved catalytic cysteine and histidine residues are identified.

**Fig S2** Purification of 6His::CNF1. **(A)** SDS PAGE gel of NiNTA purification steps of 6His::CNF1. (1) Pre-induction, (2) Post-induction, (3) Insoluble lysate fraction, (4) Soluble lysate fraction, (5) Post-resin incubation flowthrough, (6) Wash flowthrough, (7) 5mM ATP wash flowthrough (8) Post-wash resin, (9-12) Elutions 1-4, (13) Post-elution resin. **(B)** SDS PAGE gel of FPLC gel filtration fractions. (1 & 22) Input, (2-21) Fractions 25-45. **(C)** Spectrogram of FPLC gel filtration of combined NiNTA elutions of 6His::CNF1. **(D)** SDS PAGE gel showing final stock of purified 6His::CNF1 and 6His::CNF1 C866A purified by identical means.

**Fig S3** Comparison of CAB2 and CAB2Δ*vopC*. Each data point represents an average OD_600_ measurement derived from three technical replicates. No significantly differences in average OD_600_ were observed between the assayed strains.

